# Hematobiochemical Alterations and Pathological Lesions Induced by Fasciolosis in Slaughtered Cattle at Gondar ELFORA Abattoir, Ethiopia

**DOI:** 10.1101/2024.08.10.607441

**Authors:** Abraham Belete Temesgen, Tadegegn Mitiku, Mastewal Birhan, Mersha Chanie Kebede, Mohammed Yesuf, Muluken Yayeh Mekonnen, Moges Maru Alemayehu, Birhan Anagaw Malede, Abdo Megra Geda, Aregash Wendimu Tumebo, Kefale Ambachew Shiferaw, Zerihun Getie Wassie, Genetu Kassahun Berie, Bemrew Admassu Mengistu, Melaku Getahun Feleke, Fikadu Edenshaw, Mulusew Tesfaye Yitie, Gashaw Enbiyale Kasse, Elias Melkamu Tsehay, Samuel Atalay Shiferaw

## Abstract

Fasciolosis is a neglected zoonotic parasitic disease caused by *Fasciola hepatica* and *Fasciola gigantica*. It is a serious public health and veterinary concern, with significant consequences for both human and livestock populations. An abattoir-based cross-sectional study was conducted from January to September 2023 at Gondar ELFORA Abattoir, Northwest Ethiopia, to assess hematobiochemical alterations and pathological lesions induced by fasciolosis in slaughtered cattle.The study included one hundred apparently healthy male local-breed cattle, divided into two groups of fifty: one infected and one non-infected. Cattle were selected using a purposive sampling technique. Infected cattle showed significantly lower mean values for hemoglobin (HGB), hematocrit (HCT), mean corpuscular volume (MCV), total erythrocyte count (TEC), lymphocytes, monocytes, total protein, albumin, and glucose compared to non-infected cattle. Conversely, they had higher mean values for mean corpuscular hemoglobin concentration (MCHC), total leukocyte count (TLC), neutrophils, eosinophils, aspartate aminotransferase (AST), alanine aminotransferase (ALT), and alkaline phosphatase (ALP). Basophil levels were similar in both groups. Liver alterations observed in acute fasciolosis included hepatomegaly with rounded edges and the presence of juvenile flukes within the parenchyma, while in chronic fasciolosis, the liver appeared smaller, firm, with a corrugated capsule and dilated bile ducts containing twisted flukes. Microscopically, acute fasciolosis showed eosinophil infiltration, hemosiderin pigmentation, and congestion around the central vein and sinusoids, whereas chronic fasciolosis showed fibrosis, bile duct proliferation, and metaplasia of epithelial cells from columnar to cuboidal. The observed findings indicate a severe hepatic and systemic disease process driven by parasitic infection, resulting in significant compromise of the cattle’s health. Therefore, regular screening and effective deworming are essential to control bovine fasciolosis, especially in high-risk abattoirs. Hematology and biochemical tests should be part of routine diagnosis for early detection and liver function assessment, while histopathology confirms the infection stage. Enhancing stakeholder awareness and training is vital, and further research on seasonal patterns, risk factors, and drug resistance is needed to improve control strategies.

## 1. Introduction

Fasciolosis is a neglected zoonotic parasitic disease caused by *Fasciola hepatica* and *Fasciola gigantica*. It is a serious public health and veterinary concern, with significant consequences for both human and livestock populations(Zeng et al. 2023). *Fasciola*, often known as the liver fluke, causes both acute and chronic inflammation of the liver. Submandibular edema, cirrhosis, weight loss, anemia, hypoproteinemia, and anorexia are some of the subsequent symptoms. The larger species, *F. gigantica*, suppresses the host’s immune system, enhancing its pathogenicity and fatality (Yusuf et al. 2016; Ashoor and Wakid 2023). All mammals can be infected by these species, but cattle are especially vulnerable to these parasites (Beesley et al. 2018).

Acute fasciolosis occurs when immature flukes cause significant liver damage, hemorrhage, and fibrinous exudates before migrating to the bile ducts. During the chronic phase, the flukes settle in the bile ducts, causing anemia, hypoproteinemia, and cirrhosis. The bile ducts are typically swollen, cystic, and calcified, especially in cattle (Costa et al. 2022). Hematological assessments are essential for diagnosing systemic diseases (Roland et al. 2014). Fasciolosis disrupts liver metabolic activities by damaging hepatocytes, causing changes in biochemical indicators such as glucose, liver enzymes, and serum proteins (Gattani et al. 2018; Yesuf et al. 2020). Evaluating these alterations is vital for assessing hepatic damage caused by the disease (Abd Ellah et al. 2014; Gattani et al. 2018; Yesuf et al. 2020; Brahmbhatt et al. 2021).

Abattoirs provide valuable epidemiological data on diseases through meat inspection procedures, including antemortem and postmortem examinations. While backyard slaughter is common in Ethiopia (Mesfin and Mekonnen 2014), recent studies have focused on gross pathological examinations to characterize lesions and quantify economic losses due to fasciolosis (Dechasa et al. 2012; Meaza et al. 2017). However, integrated studies linking hematobiochemical alterations with pathological lesions in bovine fasciolosis remain limited (Kitila and Megersa 2014). Therefore, this study aimed to characterize the gross and microscopic lesions in *Fasciola*-infected cattle and compare their hematobiochemical profiles with those of non-infected cattle.

## 2. Materials and Methods

### 2.1. Study area and animal

The study was conducted from January to September 2023 at the Gondar ELFORA abattoir, a privately owned facility that handles sheep, goats, and cattle in Gondar, the capital of the Central Gondar administrative zone in the Amhara regional state (Figure 1). Gondar is located at latitude of 12° 36’ N and a longitude of 37° 28’ E, with an altitude ranging from 1,800 to 2,220 meters above sea level. It is approximately 727 km from Addis Ababa. The area receives 880-1,172 mm of annual rainfall, mostly during the wet season from September to May. The average temperature is 20.3°C, with 55.7% relative humidity (Tariku et al.2018). The abattoir primarily serves the local market and the University of Gondar, slaughtering 25-30 cattle weekly from various districts within the Central Gondar zone. Male, apparently healthy, local breed cattle brought for slaughter for human consumption were subjected to the study. These cattle were raised in an extensive farming system, transported from their origin to the abattoir by trucks, and kept in lairage for one day.

**Figure 1:**
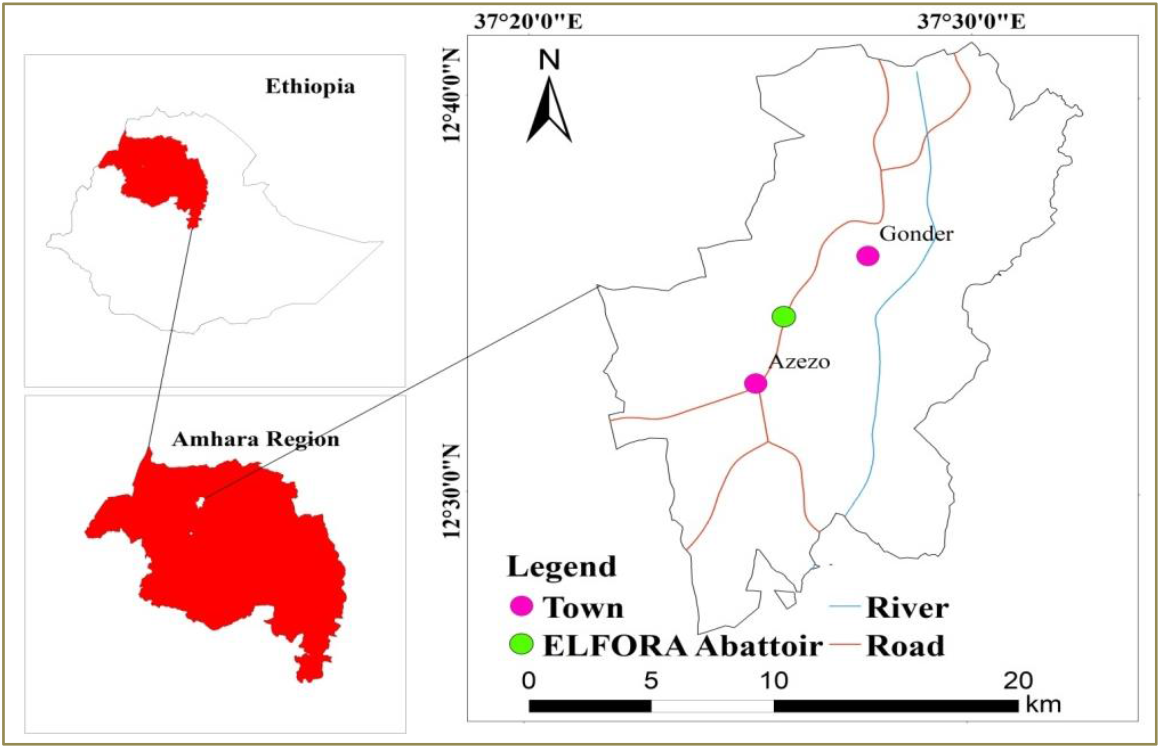
Map of the study area(ArcGIS 10.8)

### 2.2. Study design and period

An abattoir-based cross-sectional study was conducted from January to September 2023 to assess the hematobiochemical alterations and pathological lesions induced by fasciolosis in slaughtered cattle at Gondar ELFORA Abattoir.

### 2.3. Sample size and method

A total of 100 apparently healthy male local-breed cattle were purposively selected for this study and divided into two groups of 50: one infected and one non-infected. Group classification was based on the confirmed presence of *Fasciola* spp. in the liver, as determined by gross examination, which included systematic incision, palpation, and visual inspection of the liver tissue.

### 2.4. Inclusion and exclusion criteria

Blood samples collected during the pre-slaughter phase were included in the infected group only after the presence of *Fasciola* was confirmed through gross or postmortem examination of the liver. Samples from cattle in which *Fasciola* was not detected were classified in the non-infected group.

### 2.5. Sample collection and transportation

A total of 100 blood samples were collected using 5 mL EDTA-coated vacutainers for hematological analysis and 5 mL plain tubes for serum biochemical analysis. Additionally, 25 representative liver tissue samples, approximately 3 cm in length, were harvested from 50 *Fasciola*-infected cattle. Each sample included both infected tissue and adjacent healthy tissue for histopathological examination. The liver samples were preserved in 10% neutral-buffered formalin, while the blood samples were stored on ice packs during transport. Fecal samples were also collected directly from the rectum of cattle and stored in universal bottles containing 10% neutral-buffered formalin for further analysis.All specimens were subsequently transported to the Veterinary Pathology Laboratory at the University of Gondar.

### 2.6. Sample processing

#### 2.6.1. Hematobiochemical analysis

Hematology analysis was performed on blood samples treated with EDTA and stored at +4°C. A Sysmex automated hematology analyzer was used to measure parameters including hemoglobin (HGB), hematocrit (HCT), total erythrocyte count (TEC), total leukocyte count (TLC), and differential leukocyte count (DLC), as well as calculated values such as mean corpuscular volume (MCV), mean corpuscular hemoglobin (MCH), and mean corpuscular hemoglobin concentration (MCHC). For biochemical analysis, anticoagulant-free blood samples were allowed to clot and the serum was subsequently stored at -20°C. The serum concentrations of aspartate aminotransferase (AST), alanine aminotransferase (ALT), alkaline phosphatase (ALP), total protein, albumin, and glucose were determined with a YSI automated clinical chemistry analyzer.

#### 2.6.2. Gross examination

The liver and associated biliary structures were examined during postmortem inspection, both before and after dissection. The procedure included visual inspection, palpation, and incision of the liver, bile ducts, and gallbladder. Livers showing fasciolosis lesions and the presence of *Fasciola* species were selected for histopathological examination. Additionally, bile samples were collected for further analysis. *Fasciola* specimens from the bile ducts and gallbladders were harvested in normal saline, following the protocol outlined by Javaregowda and Rani (2017).

#### 2.6.3. Histopathology examination

Twenty-five liver tissue samples infected with *Fasciola* spp. were fixed in 10% neutral-buffered formalin, dehydrated in ascending grades of ethanol, cleared with xylene, impregnated, and embedded in paraffin wax, and then sectioned at 5 μm using a microtome. The sections were stained with hematoxylin and eosin (H&E), mounted with dibutyl phthalate xylene (DPX), and coverslipped. The slides were then properly dried and examined under a light microscope at objective magnifications ranging from 10x to 100x, as described by Al-Mahmood and Al-Sabaawy (2019).

### 2.7. Analysis of data and statistical methods

The data obtained were arranged, checked, coded, and entered into an Excel spreadsheet (Microsoft Office Excel 2010). They were then analyzed using STATA version 16. Significant differences between the hematological and biochemical parameters of the *Fasciola*-infected and non-infected samples were determined by t-test. Differences were considered significant when P<0.05, and the results were expressed as means ± SE. Images depicting both microscopic and gross lesions were utilized to illustrate descriptive statistics.

## 3. Results

### 3.1. Hematobiochemical findings

The mean hematology and serum biochemical parameters of *Fasciola*-infected and non-infected cattle are compared in Tables 1 and 2. Infected cattle showed lower mean values for hemoglobin (HGB), hematocrit (HCT), mean corpuscular volume (MCV), total erythrocyte count (TEC), lymphocytes, monocytes, total protein, albumin, and glucose. Conversely, they had higher mean values for mean corpuscular hemoglobin concentration (MCHC), total leukocyte count (TLC), neutrophils, eosinophils, aspartate aminotransferase (AST), alanine aminotransferase (ALT), and alkaline phosphatase (ALP). Basophil levels were similar in both groups. The parameters of non-infected cattle were within reference ranges. Significant differences (P<0.05) were observed in HGB, HCT, MCV, TEC, TLC, lymphocytes, monocytes, neutrophils, eosinophil, AST, ALT, ALP, total protein, albumin, and glucose between infected and non-infected cattle, while mean corpuscular hemoglobin (MCH) and MCHC showed no significant differences.

**Table 1:** Hematological parameters of *Fasciola-*infectedand non-infected cattle.

| Parameters | Mean ± SE | | Range | t-value | p-value |
| --- | --- | --- | --- | --- | --- |
|  | Infected<br>(n=50) | Non-infected<br>(n=50) |  |  |  |
| HGB(g/dL)* | 4.76 ± 0.18 | 9.76 ± 0.19 | 8-14 | 20.34 | 0.000 |
| HCT(%)* | 15.96 ± 0.62 | 34.98 ± 0.81 | 24-42 | 19.52 | 0.000 |
| MCV(fL)* | 49.10 ± 2.23 | 56.42 ± 1.59 | 40-60 | 2.87 | 0.005 |
| MCH(pg)<sup>NS</sup> | 15.05 ± 0.88 | 15.78 ± 0.44 | 11-17 | 0.74 | 0.457 |
| MCHC (%)<sup>NS</sup> | 33.18 ± 2.23 | 28.89 ± 1.07 | 28-36 | 2.00 | 0.0509 |
| TEC ($\times 10^6/\mu\text{L}$)* | 3.37 ± 0.10 | 6.29 ± 0.11 | 5-10 | 18.17 | 0.000 |
| TLC ($\times 10^3/\mu\text{L}$)* | 14.35 ± 0.18 | 6.41 ± 0.22 | 4-12 | 29.09 | 0.000 |
| LYM ($\times 10^3/\mu\text{L}$)* | 1.66 ± 0.05 | 3.17 ± 0.17 | 2.5-7.5 | 8.79 | 0.000 |
| MON ($\times 10^3/\mu\text{L}$)* | 0.019 ± 0.001 | 0.79 ± 0.08 | 0.02-0.85 | 9.60 | 0.000 |
| NEU ($\times 10^3/\mu\text{L}$)* | 4.35 ± 0.21 | 2.52 ± 0.14 | 0.6-4 | 8.33 | 0.000 |
| EOS ($\times 10^3/\mu\text{L}$)* | 3.25 ± 0.135 | 0.53 ± 0.037 | 0-2.4 | 18.19 | 0.000 |
| BAS ($\times 10^3/\mu\text{L}$)<sup>NC</sup> | 0 ± 0 | 0 ± 0 | 0-0.2 | - | - |

Note: Reference range adopted from Sirois and Hendrix (2015). Abbreviations: HGB = Hemoglobin, HCT = Hematocrit, TEC = Total erythrocyte count, MCV = Mean corpuscular volume, MCH = Mean corpuscular hemoglobin, MCHC = Mean corpuscular hemoglobin concentration, TLC = Total leukocyte count, LYM = Lymphocytes, MON = Monocytes, NEU = Neutrophils, EOS = Eosinophils, BAS = Basophils. \*Significant at $P < 0.05$ . NS = Non-significant at $P < 0.05$ . NC = No change between groups.

**Table 2:** Biochemical parameters of *Fasciola*-infected and non-infected cattle.

| Parameters | Mean ± SE | | Range | t-value | p-value |
| --- | --- | --- | --- | --- | --- |
|  | Infected<br>(n=50) | Non-infected<br>(n=50) |  |  |  |
| AST(U/L)* | 244.78 ± 5.63 | 108.52 ± 3.31 | 46-176 | 20.37 | 0.000 |
| ALT(U/L)* | 61.12 ± 1.96 | 21.46 ± 0.86 | 5-35 | 17.43 | 0.000 |
| ALP (U/L)* | 195.7 ± 3.11 | 69.14 ± 5.02 | 18-153 | 23.90 | 0.000 |
| TP (g/dL)* | 3.92 ± 0.17 | 6.51 ± 0.057 | 6-7.5 | 13.84 | 0.000 |
| Albumin (g/dL)* | 2.43 ± 0.086 | 3.79 ± 0 .035 | 3.4-4.3 | 14.03 | 0.000 |
| Glucose (g/dL)* | 25.14 ± 0.71 | 55.74 ± 1.42 | 37-79 | 19.22 | 0.000 |

Note: Reference range adopted from Sirois and Hendrix (2015). Abbreviations: ALT = Alanine aminotransferase, AST = Aspartate aminotransferase, ALP = Alkaline phosphatase, TP = Total protein.
\*Significant at $P<0.05$ .

### 3.2. Fecal and bile analysis

The eggs of *Fasciola* species detected in cattle fecal and bile samples in this study are presented in Figure 2. The identification of these eggs was based on the morphological characteristics outlined by Yakhchali and Bahramnejad (2015), Karimov et al. (2021), and Ngazizah et al. (2023). These eggs are characteristically ovoid in shape, with a robust, yellowish-brown shell, and often feature an operculum at one end.

**Figure 2:**
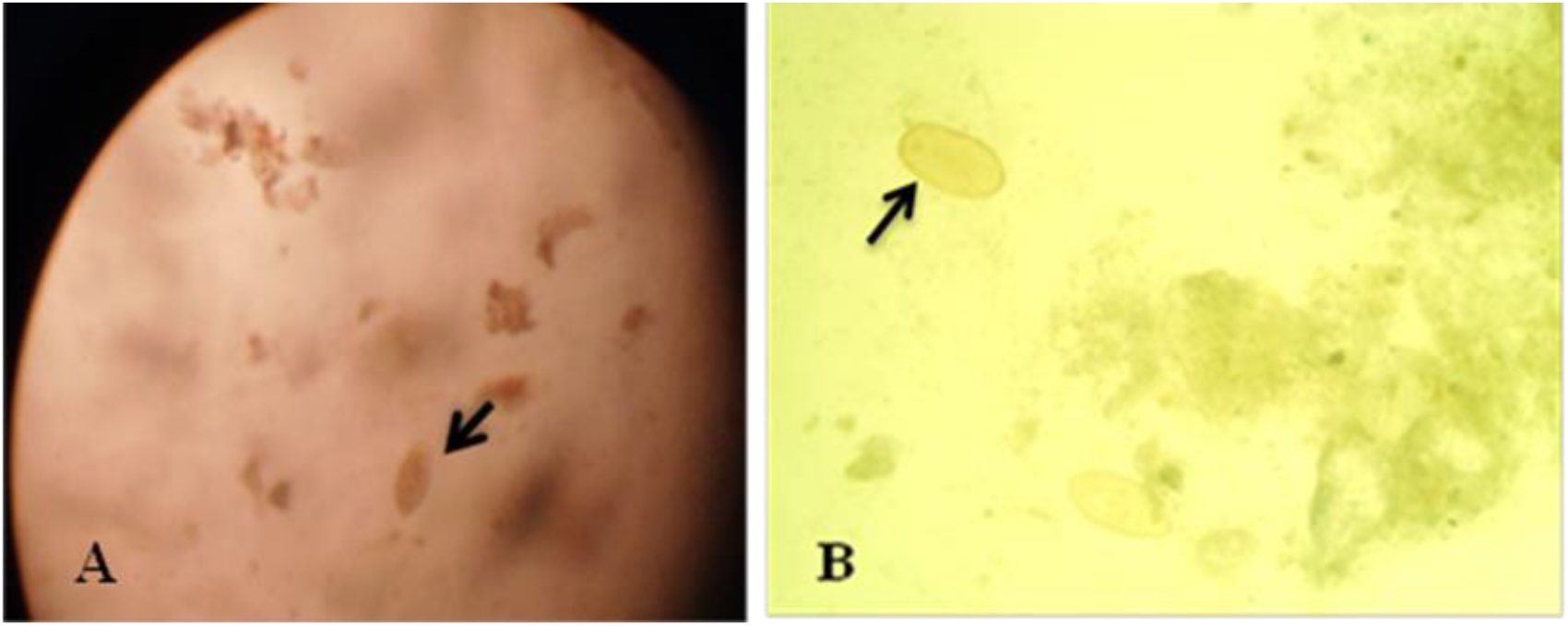
Eggs of *Fasciola* species in chronic fasciolosis cases: yellowish eggs in fecal (A) and bile smear (B).

### 3.3. Gross and microscopic lesions

#### 3.3.1. Gross lesions

In acute fasciolosis, the livers exhibited hepatomegaly with rounded edges due to parenchymal inflammation, and pinpoint hemorrhages were present on the parietal surface. Some areas of the parenchyma were pale due to necrosis. The capsule was firm, rough, and thick with whitish or reddish discoloration within the parenchyma. A juvenile fluke was observed in the cut section (Figure 3A). There was also congestion and engorgement of the gallbladder with bile, and upon dissection, a blackish-brown exudate was found. The mucosa was normal, and no flukes or ova/eggs were present (Figure 3B).

**Figure 3:**
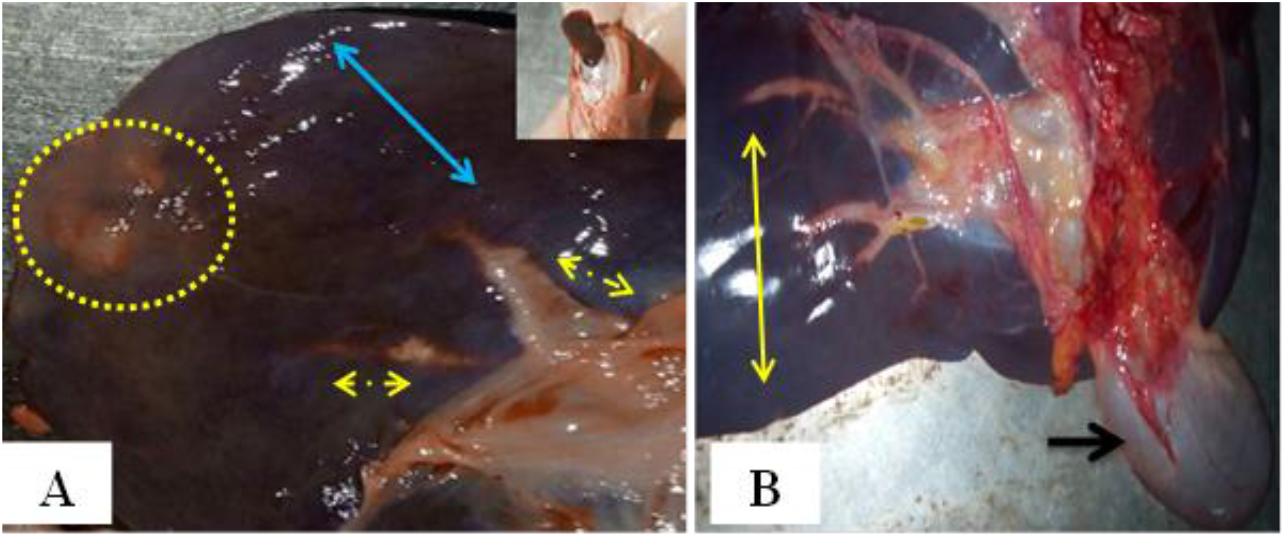
Gross lesions of acute fasciolosis: (A) Hepatomegaly with rounded edges and pinpoint hemorrhages (blue arrows) on the parietal surface, paleness in some areas (dotted arrows), a firm, rough, and thick capsule with whitish or reddish discoloration within the parenchyma (dotted circle), and a juvenile fluke (inset) seen at the cut section. (B) Congestion (yellow arrows) and engorgement of the gallbladder with bile (black arrow).

In chronic fasciolosis,the liver was small in size and firm in consistency with a corrugated capsule (Figure 4A). The bile duct was engorged (Figure 4B). Upon dissection, numerous mature and adult twisted flukes were found along with blackish-brown exudate, giving the liver a pipe stem appearance. The gallbladder was reduced in size, and its mucosa was thickened and contained flukes and ova/eggs (Figure 4C). The flukes collected from the bile ducts and gallbladder were identified as *Fasciola hepatica* and *Fasciola gigantica* (Figure 4D).

**Figure 4:**
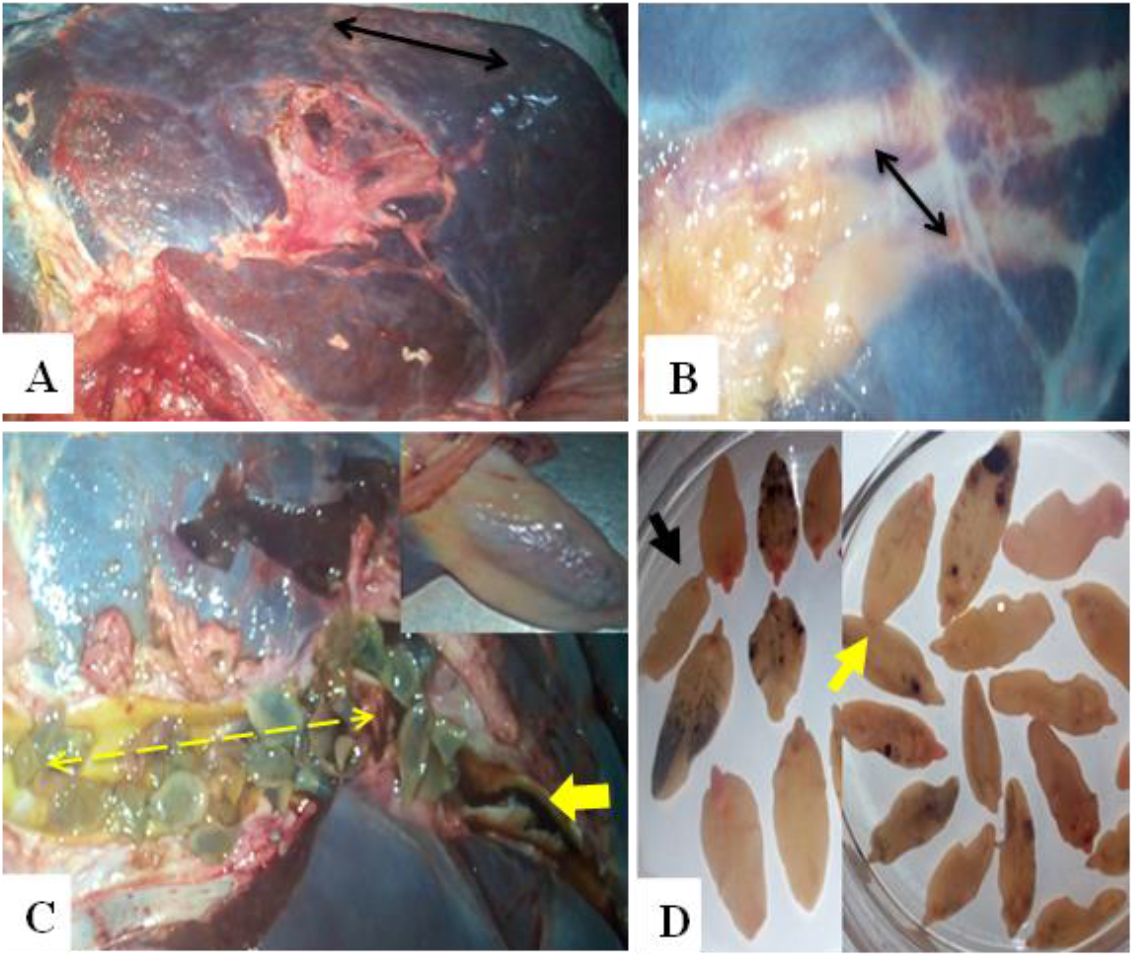
Gross lesions of chronic fasciolosis: (A) the liver was small in size and firm in consistency with a corrugated capsule (arrows). (B) Engorged bile ducts (arrows). (C) Numerous mature and immature twisted flukes (lined arrows) along with blackish-brown exudate (yellow arrow), and the gallbladder (inset) was decreased in size. (D) Identification of *F. hepatica* (black arrow) and *F. gigantica* (yellow arrow).

#### 3.3.2. Microscopic lesions

Inacute fasciolosis, excessive eosinophil infiltration was observed in migratory tracts (Figure 5A), along with congestion around the central vein due to dilation of the central vein and sinusoids engorged with a large number of RBCs (Figure 5B). Hemosiderin pigmentation was also noted (Figure 5C). A necrotic area surrounded by degenerating swollen hepatocytes and inflammatory cells, with RBC engorgement, was identified (Figure 5D). Coagulative necrosis was observed in the migratory tracts, surrounded by large clear vacuoles, and along with pyknosis, karyolysis, and karyorrhexis within the cytoplasm (Figure 5E). Swollen hepatocytes with collapsed cytoplasm, hyperplasia of fibrocytes, and congested blood vessels at the portal area were also recorded (Figure 5F).

**Figure 5:**
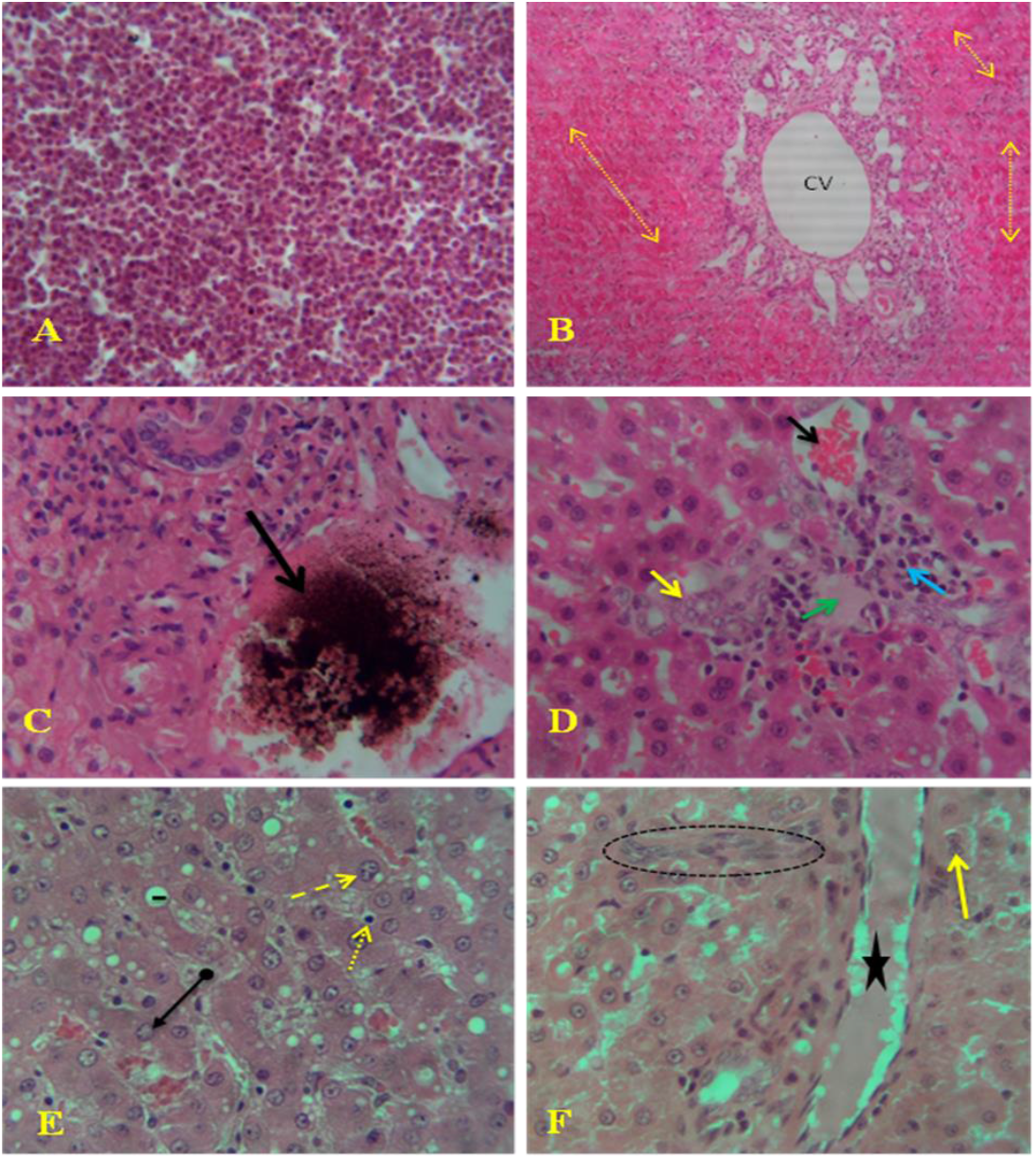
Microscopic lesions of acute fasciolosis: (A) Excessive eosinophil infiltration in migratory tracts. (B) Congestion around the central vein (CV) with engorgement of red blood cells (dotted arrows), 10X magnification. (C) Hemosiderin pigmentation (arrow). (D) Necrotic area (green arrow) with engorgement of red blood cells (black arrow), surrounded by degenerating swollen hepatocytes (yellow arrows) and inflammatory cells (blue arrow). (E) Coagulative necrosis (arrow tail) in the migratory tracts, surrounded by a large clear vacuole (line), pyknosis (dotted arrow), karyolysis (arrowhead), and karyorrhexis (lined arrow) within the cytoplasm. (F) Swollen hepatocytes with collapsed cytoplasm (arrow), hyperplasia of fibrocytes (circle), and congested blood vessels around the portal area (star). H&E. 40X.

In chronic fasciolosis, neutrophil infiltration was observed within the sinusoidal capillaries and among necrotic hepatocytes (Figure 6A). There was proliferation of fibrous connective tissues with fibrosis and hemorrhage (Figure 6B). Fatty changes were noted along with metaplasia of columnar epithelial cells and bile duct proliferation (Figure 6C). Metaplasia of columnar to cuboidal epithelial cells, along with metaplasia of columnar epithelial cells, was also observed (Figure 6D).

**Figure 6:**
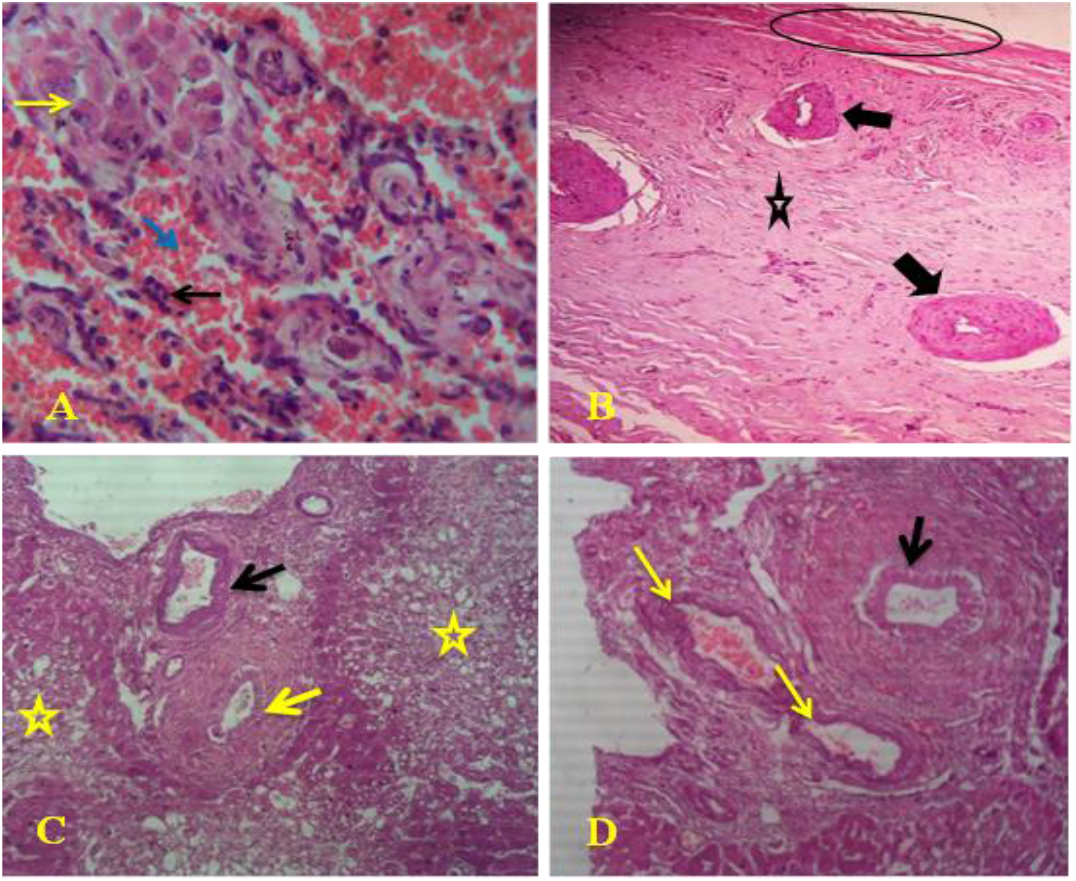
Microscopic lesions of chronic fasciolosis: (A) Neutrophil infiltration (black arrow) within sinusoidal capillaries (blue arrow) and among necrotic hepatocytes (yellow arrow). (B) Proliferation of fibrous connective tissues (arrows) with fibrosis (star) and hemorrhage (circle). (C) Fatty changes (stars) along with metaplasia of columnar epithelial cells (black arrow) and bile duct proliferation (yellow arrow). (D) Metaplasia of columnar to cuboidal epithelial cells (yellow arrows) and metaplasia of columnar epithelial cells (black arrow). H&E. 40X.

## 4. Discussion

The diagnosis of bovine fasciolosis was confirmed through hematobiochemical analyses, characterization of pathological lesions, and examination of fecal and bile samples. Significant reductions were observed in hemoglobin (HGB), hematocrit (HCT), mean corpuscular volume (MCV), total erythrocyte count (TEC), lymphocytes, monocytes, total protein (TP), albumin, and glucose in *Fasciola*-infected cattle. Conversely, the total leukocyte count (TLC), eosinophils, neutrophils, aspartate aminotransferase (AST), alanine aminotransferase (ALT), and alkaline phosphatase (ALP) were elevated. Basophil levels were similar in both infected and non-infected cattle. These findings are consistent with previous studies (Hossain et al. 2006; Singh et al. 2011; Abd Ellah et al. 2014; Gattani et al. 2018; Brahmbhatt et al. 2021).

Reductions in HCT, TEC, HGB, and MCV, along with an elevated MCHC, are indicative of microcytic hyperchromic anemia. This clinical manifestation is commonly attributed to blood loss resulting from hemorrhages caused by migrating immature flukes and blood-feeding adult flukes, as reported by Brahmbhatt et al. (2021). Similarly, Maskur et al. (2022) documented significant reductions in HCT, HGB, and TEC in association with liver fluke infestations and subsequent hepatic damage. Comparable hematological changes have been observed in several studies, including those by Egbu et al. (2013), Van Wyk et al. (2013), Gattani et al. (2018), Ngetich (2019), and Brahmbhatt et al. (2021).

Moreover, significant reductions in HCT, TEC, and HGB have been documented in *Fasciola*-infected sheep, as reported by Ahmed et al. (2006), Matanović et al. (2007), and Yesuf et al. (2020). A pronounced decrease in mean corpuscular volume (MCV), along with a non-significant reduction in MCH and an elevated MCHC, aligns with the findings of Taimur et al. (1993), who observed similar trends in *Fasciola*-infected cattle. In contrast, Yesuf et al. (2020) noted significant reductions in MCV, MCH, and MCHC in infected sheep, while Coppo et al. (2011) identified a non-significant reduction in MCV, with concurrent elevations in MCH and MCHC in infected cattle. These variations may be attributable to ecological and nutritional factors that influence the hematological profiles of different host species, as discussed by Yesuf et al. (2020).

Elevations in TLC, eosinophils, and neutrophils in liver fluke-infected cattle are indicative of leukocytosis, eosinophilia, and neutrophilia, while reductions in lymphocytes and monocytes suggest lymphocytopenia and monocytopenia, respectively. These hematological alterations are consistent with findings from several studies (Taimur et al. 1993; Egbu et al. 2013; Brahmbhatt et al. 2021). Ganguly et al. (2016) reported significant leukocytosis, neutropenia, and lymphocytopenia, with non-significant eosinophilia and monocytosis in *Fasciola*-infected sheep. Similarly, Yesuf et al. (2020) observed significant leukocytosis and eosinophilia, along with non-significant neutropenia, lymphocytosis, and monocytopenia in sheep. El-Aziem Hashem and Mohamed (2017) also documented significant increases in leukocytosis, eosinophilia, and neutrophilia in *Fasciola*-infected cattle. Elevation in total leukocyte count (TLC) may reflect a physiological response aimed at combating liver fluke infestation (Maskur et al. 2022).

Eosinophilia is commonly associated with immune defense mechanisms against parasitic infections (Yesuf et al. 2020), possibly involving the release of cytotoxic molecules from the surface of the parasites (Piedrafita et al. 2001). Neutrophilia, a frequent finding, may be linked to secondary bacterial infections resulting from fluke migration through the biliary parenchyma (Taimur et al. 1993) or the phagocytic activity of neutrophils involved in degrading the parasite’s cuticle. Furthermore, lymphocytopenia observed in these infections may be related to immune responses, with a concurrent increase in neutrophil counts reflecting the body’s heightened inflammatory response (Katre et al. 2020).

Elevations in AST, ALT, and ALP levels, along with reductions in TP, albumin, and glucose levels, observed in *Fasciola*-infected cattle, are indicative of liver damage resulting from fluke migration. These findings are consistent with the studies of Hodžić et al. (2013) and Brahmbhatt et al. (2021), which emphasize that increased AST and ALT levels are reflective of inflammatory responses, fibrosis, and necrosis induced by the parasitic infestation. Elevated ALP levels, as noted by Coppo et al. (2011) and Brahmbhatt et al. (2021), suggest cholestasis due to bile duct obstruction caused by migrating flukes.Reductions in protein and albumin levels, indicative of hypoproteinemia and hypoalbuminemia, are likely a consequence of hepatocellular damage induced by the flukes, which impair the liver’s ability to synthesize proteins. Furthermore, the decreased glucose levels, indicative of hypoglycemia, are believed to result from disrupted hepatic glycogenesis due to fluke migration, aligning with the findings of Hossain et al. (2006), Gattani et al. (2018), and Yesuf et al. (2020).

In acute fasciolosis, macroscopic changes included hepatomegaly, rounded liver edges with parenchymal inflammation, pinpoint hemorrhages, congestion, pale areas indicative of necrosis, and a firm, rough capsule with discoloration within the parenchyma. Juvenile flukes were observed, consistent with previous studies across various livestock species (Ahmedullah et al. 2007; Sayed et al. 2008; Metwally et al. 2009; Borai et al. 2013; Okoye et al. 2015; Kitila and Megersa 2014; Mohamed et al. 2021). Additionally, an engorged gallbladder filled with bile, along with the presence of a blackish-brown exudate upon dissection, aligns with findings from previous studies (Adrien et al. 2013; Islam et al. 2016). The absence of fluke eggs in both the gallbladder and fecal samples suggests an early stage of infection. These observations are consistent with reports indicating that variations in gallbladder and bile duct abnormalities may be influenced by breed differences (Simpson et al. 2009; Lamb et al. 2021).

In chronic fasciolosis, the liver appeared small and firm, with a corrugated capsule. The bile duct was engorged, containing both immature and adult flukes, along with a blackish-brown exudate, giving the liver a characteristic pipe-stem appearance. These findings are consistent with previous studies across various livestock species (Borai et al. 2013; Kitila and Megersa 2014; Salmo et al. 2014; Islam et al. 2016; Mohamed et al. 2021). The gallbladder exhibited thickened, edematous mucosa containing flukes and eggs, contrasting with Adrien et al. (2013), who reported thickened mucosa without mentioning size reduction. Reduced gallbladder size may be attributed to bile duct obstruction by flukes, impairing bile synthesis. The presence of fluke eggs in both bile and fecal samples might be due to the chronic stage of fasciolosis, when adult *Fasciola* species living in the bile ducts start laying eggs, which are then passed into the intestine and excreted in the feces (Inoue et al. 2007; Nu et al. 2022).The identification of *F*.gigantica and *F. hepatica* in the gallbladder and bile duct was based on their characteristic morphology (Giovanoli Evack et al. 2020; Sumruayphol et al. 2020).

Microscopic changes observed in acute fasciolosis included eosinophil infiltration, sinusoidal dilation with RBC engorgement, hepatocyte necrosis, and hemosiderin deposition. In chronic fasciolosis, histopathological examinations consistently revealed several features across different species infected with *Fasciola*, including neutrophil infiltration in sinusoidal capillaries, necrotic hepatocytes, and fibrous tissue proliferation in bile ducts, fibrosis, fatty changes, and epithelial cell metaplasia. These observations are supported by other studies (Borai et al. 2013; Salmo et al. 2014; Al-Mahmood and Al- Sabaawy 2019; Mohamed et al. 2021; Ashoor and Wakid 2023).

## 5. Conclusion

This study confirmed that bovine fasciolosis induces significant hematological, biochemical, gross pathological, and histopathological alterations. Cattle infected with *Fasciola* developed a complex pathological condition characterized by anemia with abnormal red blood cell morphology, severe liver injury including hepatocellular necrosis, disrupted bile flow causing cholestasis and biliary obstruction, compensatory bile duct hyperplasia, and metabolic disturbances such as glycogen depletion and impaired protein synthesis. The observed findings indicate a severe hepatic and systemic disease process driven by parasitic infection, resulting in significant compromise of the cattle’s health. Therefore, regular screening and effective deworming are essential to control bovine fasciolosis, especially in high-risk abattoirs. Hematology and biochemical tests should be part of routine diagnosis for early detection and liver function assessment, while histopathology confirms the infection stage. Enhancing stakeholder awareness and training is vital, and further research on seasonal patterns, risk factors, and drug resistance is needed to improve control strategies.

## Data sharing statement

All data generated or analyzed during this study are available upon request from the corresponding author.

## Ethics statement

This study was approved by the Ethics Research Review Committee of the University of Gondar College of Veterinary Medicine and Animal Sciences (Ref. No. CVMASc/UoG/RERC/20/12/2022) on December 20, 2022. All animal procedures adhered to established ethical guidelines, and verbal informed consent was obtained from the Gondar ELFORA abattoir manager, as approved by the ethics committee, prior to sample collection.

## Acknowledgments

The authors thank the University of Gondar, College of Veterinary Medicine and Animal Sciences, for its support in all aspects, as well as the staff of the Gondar ELFORA Abattoir for their invaluable assistance during the sample collection process.

## Author contributions

ABT: writing the original draft, reviewing and editing, funding acquisition, resourcing, methodology, visualization, validation, investigation, conceptualization, formal analysis, and data curation. TM and MB: supervision, visualization and validation. MCK, MY, MYM, and MMA: visualization, validation, methodology and review editing. BAM, AMG, AWT, KAS, ZGW, GKB, BAM, MGF, FE, MTY, GEK, EMT and SAS: resourcing, review editing, visualization and validation.

